# Evidence for the agricultural origin of antimicrobial resistance in a fungal pathogen of humans

**DOI:** 10.1101/2020.05.24.113787

**Authors:** S. Earl Kang, Leilani G. Sumabat, Tina Melie, Brandon Mangum, Michelle Momany, Marin T. Brewer

## Abstract

Resistance to clinical antimicrobials is an urgent problem, reducing our ability to combat deadly pathogens of humans. Azole antimicrobials target ergosterol synthesis and are highly effective against fungal pathogens of both humans and plants leading to their widespread use in clinical and agricultural settings^1,2^. The fungus *Aspergillus fumigatus* causes 300,000 life-threatening infections in susceptible human hosts annually and azoles are the most effective treatment^3^. Resistance to clinical azole antifungals has become a major problem in Europe and India over the last decade, where identical mutations in *cyp51A*, an ergosterol biosynthetic gene, have been found in strains from both clinical and agricultural settings^4^. Shared *cyp51A* genotypes suggest that clinical azole resistance might have had an agricultural origin; however, until now, independent origins of clinical and agricultural mutations could not be ruled out. Here we show that azole-resistant isolates of *A. fumigatus* from clinical and agricultural settings also carry mutations conferring resistance to quinone outside inhibitor (QoI) fungicides, which are used exclusively in agricultural settings. This is the first report of a clear marker for the agricultural origin of resistance to a clinical antifungal. We anticipate that our work will increase the understanding of interactions between pathogens of plants and pathogens of humans.

## Main

Fungi are important pathogens of humans, causing over 1.5 million deaths annually^5^. Fungi are also important pathogens of plants, causing crop losses of 20% and postharvest losses of 10%^6^. The filamentous fungus *Aspergillus fumigatus* is a saprobe found in a variety of environments including soil, compost, and decaying plant material; however, in immunocompromised individuals it can cause the devastating disease aspergillosis. Azoles are often used as the first line of defense against aspergillosis. During the last decade Europe and Asia have seen an alarming increase in azole-resistant *A. fumigatus* in the clinic and azole resistance is now present on 6 continents^7^. Though some resistance has been associated with long-term azole therapy in patients with chronic infections, at least two-thirds of patients with azole-resistant *A. fumigatus* infections have not previously undergone azole therapy^8,9^. The environmental use of azoles has been proposed to be the driving force for the majority of clinical resistance in *A. fumigatus* with several studies suggesting that most azole-resistant isolates originated from widespread agricultural use of azoles to combat plant-pathogenic fungi^10,11^.

The same mutations in *cyp51A* – which encodes the ergosterol biosynthetic protein targeted by azoles – have been reported in strains from both clinical and agricultural settings in Europe, Asia, Africa, and the Middle East^4,7,9,12^. Several point mutations and tandem repeats (TR) within the promoter, including TR_34_/L98H and TR_46_/Y121F/T289A, are commonly associated with azole resistance in environmental isolates. Isolates with the TR_34_/L98H and TR_46_/Y121F/T289A alleles show high levels of resistance to multiple azole drugs (pan-azole resistance) and patients infected with these isolates have higher rates of treatment failure and death^13^. Though the presence of TR_34_/L98H and TR_46_/Y121F/T289A alleles in both agricultural and clinical isolates suggested that azole-resistant clinical strains of *A. fumigatus* might have had an agricultural origin, independent evolution of these mutations in both settings has not been previously excluded.

In the Unites States (U.S.), azole-resistant *A. fumigatus* strains with TR_34_/L98H and TR_46_/Y121F/T289A alleles have been recently reported in patients^14,15^. Additionally, azole-resistant *A. fumigatus* isolates with TR_34_/L98H alleles were detected in a peanut field in Georgia^16^. Otherwise, very little information about the frequency of azole-resistant *A. fumigatus* in agricultural settings in the U.S. is available. To investigate the prevalence of azole-resistant *A. fumigatus* in agricultural environments in the U.S., we collected soil and plant debris from 56 sites across Georgia and Florida, including 53 sites with a history of azole fungicide use, two organic sites with no fungicide use for at least 10 years, and one compost pile of unknown history (Supplementary Table S1). We recovered 700 isolates of *A. fumigatus* from soil, plant debris, and compost. Isolates were screened for sensitivity to the azole fungicide tebuconazole (TEB) that has widespread use in agriculture. Of the 700 isolates collected, 123 (17.6%) grew on solid medium amended with 3 μg/ml TEB. None of the isolates collected from the two organic sites grew on TEB-amended plates.

Minimal Inhibitory Concentrations (MIC) for TEB, itraconazole (ITC), voriconazole (VOR), and posaconazole (POS) were determined by broth dilution assays for the 123 isolates that grew on TEB-amended solid medium, and for 49 isolates from the same sites that that did not grow on the amended medium. MIC ranged from 0.5 to > 16 μg/ml for TEB, 0.5 to 2 μg/ml for ITC, 0.125 to > 16 μg/ml for VOR, and 0.06 to 1 μg/ml for POS. Recommended clinical breakpoints of antifungal resistance for *A. fumigatus*^17^ are > 2 μg/ml for ITC and VOR and > 0.25 μg/ml for POS; however, EUCAST notes there is uncertainty regarding the cutoff values for POS and some data suggest the cutoff value of > 1 μg/ml, which we use here, may be more relevant. Although many of the isolates showed low levels of azole resistance, only 12 of the 123 isolates were highly resistant at clinically relevant levels (Supplementary Tables S1 and S2). The 12 isolates exhibited high levels of resistance to both TEB and VOR with MIC ≥ 16 μg/ml, and decreased sensitivity to ITC and POS (Table 1), showing they are pan-azole resistant.

**TABLE 1.**
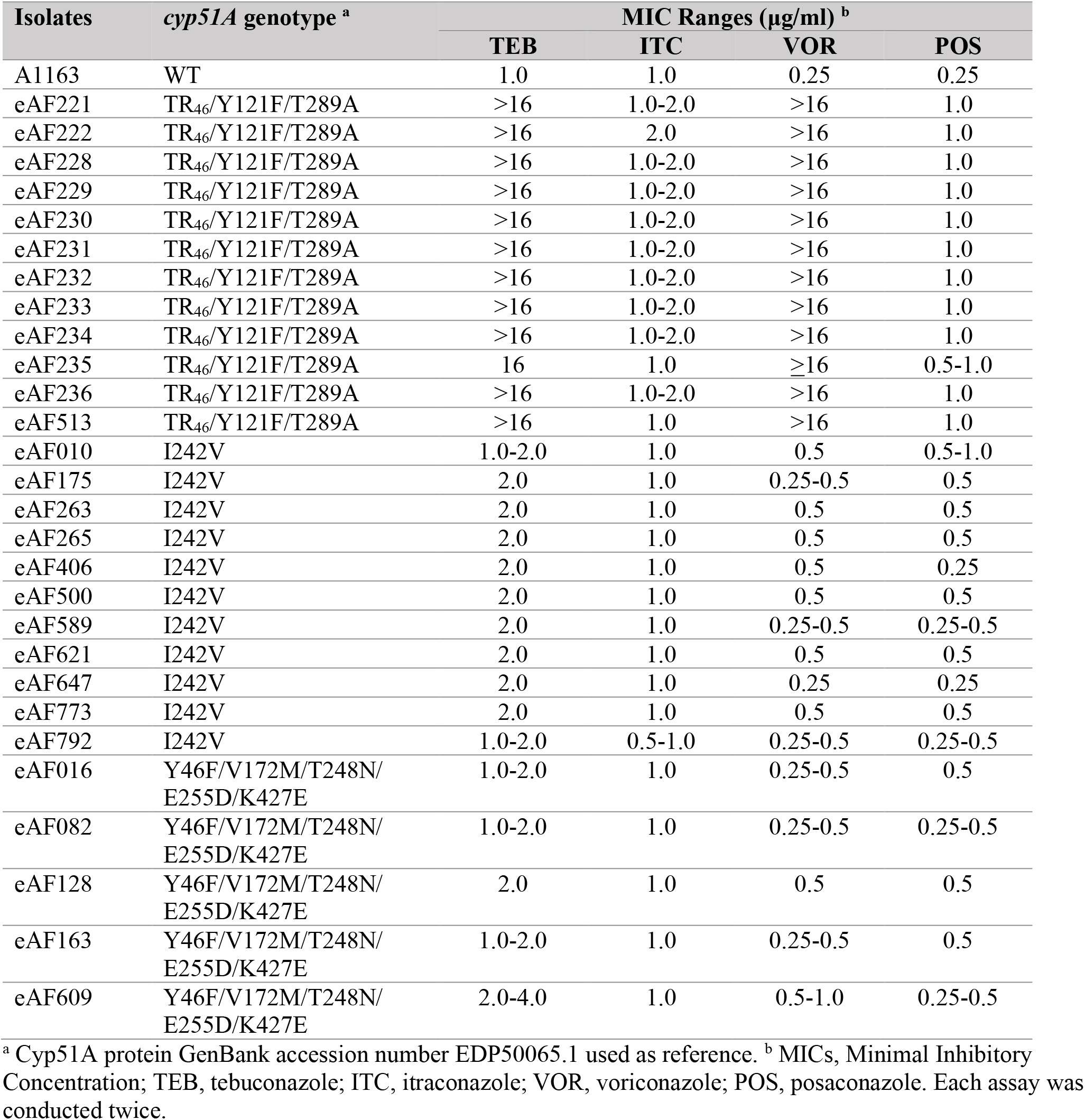
Azole-susceptibility of environmental *A. fumigatus* isolates with non-synonymous *cyp51A* mutations.

To determine whether mutations in *cyp51A* were associated with azole resistance, we sequenced 1,286 bp of *cyp51A*, including the promoter and downstream regions, for 123 isolates that grew on TEB-amended medium and for 49 TEB-sensitive isolates from the same sites. The 12 pan-azole-resistant isolates had the TR_46_/Y121F/T289A allele (Table 1) that underlies high levels of resistance to VOR^10,12^. We did not detect the TR_34_/L98H allele that is the most prevalent worldwide in azole-resistant environmental and clinical isolates of *A. fumigatus*. Failure to detect the TR_34_/L98H allele does not necessarily mean this set of mutations is absent in the areas we sampled, but more likely reflects our initial screen for resistance with TEB. TEB is an azole with a similar structure to VOR, and resistance to TEB has previously been associated with the TR_46_/Y121F/T289A mutations^18^. Eleven of the isolates sequenced had the I242V mutation in Cyp51A and 5 had the Y46F/V172M/T248N/E255D/K427E mutations found in the reference isolate Af293. These 16 isolates with non-synonymous mutations in *cyp51A,* but without tandem repeats in the promoter, had slightly elevated MIC values for TEB, VOR, and POS compared to the sensitive reference isolate A1163 (Table 1). Increased MIC values for isolates with these mutations have been described previously^19–21^; however, it is not clear if these specific mutations are the cause of increased drug resistance.

To determine the relationship of agricultural isolates from Georgia and Florida to clinical isolates from the same region, we used 9 STR*Af* markers^22^ to genotype the 168 agricultural isolates that we collected along with 48 clinical isolates collected between 2015 and 2017 by the Centers for Disease Control and Prevention^23^. None of the clinical isolates were azole resistant. Based on microsatellite data almost every isolate had a unique genotype (Supplementary Figure S1). The combined environmental and clinical *A. fumigatus* population from Georgia and Florida showed no genetic structure, except for the pan-azole-resistant agricultural isolates that had the TR_46_/Y121F/T289A allele. These pan-azole-resistant isolates comprised a single lineage and were isolated from a compost pile and pecan debris from a processing facility (Supplementary Table S1). Our results are consistent with previous studies showing that *A. fumigatus* is panmictic with little population structure either by geographic region or clinical versus agricultural setting^9,24^.

To better understand the relationship of Georgia and Florida agricultural isolates to worldwide environmental and clinical isolates, we performed whole genome sequencing on 89 strains representing all of our field sites and combined them with 69 publicly available whole genome sequences to construct a neighbor-joining tree (Figure 1; Supplementary Table S3). Clinical and environmental azole-resistant isolates (open or closed red circles) from the United States (USA), United Kingdom (UK), Spain (ESP), the Netherlands (NL), and India (IND) were distributed throughout the tree; however, the four pan-azole-resistant isolates from this study that carried *cyp51A* TR mutations (closed red circles) clustered into a well-supported clade with clinical and environmental isolates carrying *cyp51A* TR mutations from the UK, IND and NL (red branches). This clade also included azole-sensitive isolates from diverse geographic origins. Although *A. fumigatus* does not show population structure by geographic or environmental origin, our data support population structure by pan-azole resistance. The genetic relatedness of pan-azole-resistant isolates across geographic locations and environment types has been previously described^9,24^ and suggests that there is a barrier to gene flow or some other genetic predisposition in this pan-azole-resistant clade allowing *cyp51A* TR mutations to arise and/or persist.

**Figure 1.**
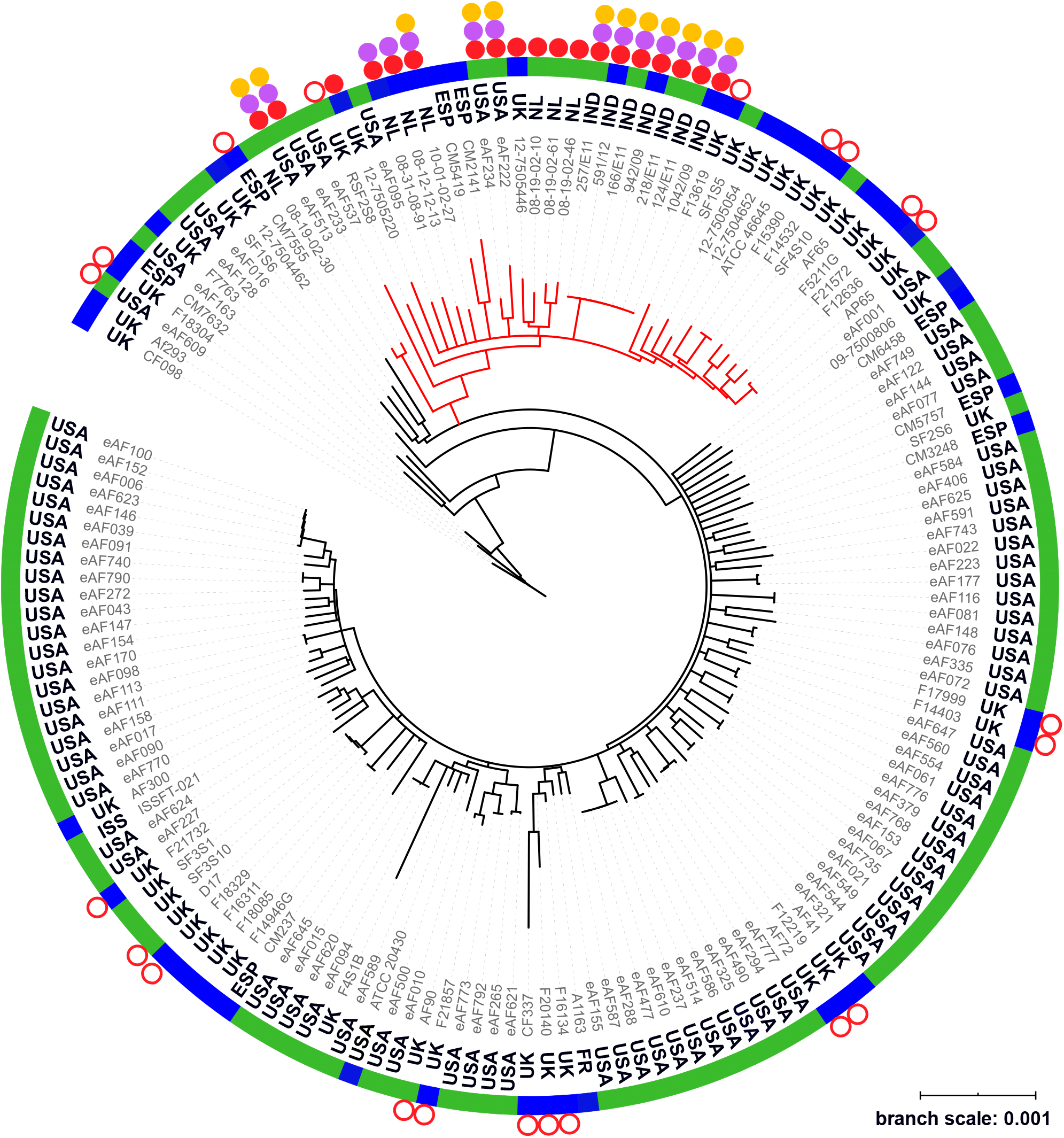
Neighbor-joining tree of environmental and clinical isolates of *Aspergillus fumigatus.* Whole genome sequences from Georgia and Florida agricultural sites (this study, eAFXXX), were analyzed along with publicly available data (Table S3). Af293 was used as the reference genome. Country of origin is listed next to each isolate (ESP=Spain, FR=France, IND=India, ISS=International Space Station, NL=Netherlands, UK=United Kingdom, USA=United States). Only branches with 100% bootstrap support based on 100 replicates are shown. Green bars indicate environmental isolates. Blue bars indicate clinical isolates. Solid red circles indicate pan-azole-resistant isolates with *cyp51A* TR mutations. Open red circles indicate azole-resistant isolates without TR mutations. Orange circles indicate isolates with *cytB* G143A mutation conferring resistance to QoI fungicides. Violet circles indicate *benA* F219Y mutation conferring resistance to MBC fungicides. Red branches indicate well-supported pan-azole-resistant clade.

To delay the evolution and spread of antifungal resistance in agricultural settings, azoles are generally applied to crops in alternation or combination with other fungicides such as the quinone outside inhibitors (QoI) and, to a lesser extent, benzimidazoles (MBC) ^25,26^. QoI fungicides target the protein encoded by the *cytochrome B* (*cytB*) gene and are used on crops, but not on patients^27^. MBC fungicides target the protein encoded by the *ß-tubulin* (*benA*) gene and were widely used in U.S. agriculture in the 1970s. MBCs are used much less in U.S. agriculture now due to resistance development, but they are used clinically as antihelminthics^28^. We reasoned that if isolates of *A. fumigatus* had acquired azole-resistance in agricultural settings, they might also have acquired resistance to these non-azole fungicides. To determine if azole-resistant isolates carried mutations conferring resistance to QoI and MBC fungicides, we searched the genomes of our agricultural isolates along with the genomes of publicly available *A. fumigatus* isolates (Supplementary Table S3). We detected the *cytB* G143A mutation known to cause QoI resistance (Figure 1, orange circles; Table 2) and/or the *benA* F219Y mutation known to cause MBC resistance (Figure 1, violet circles; Table 2) in 14 clinical and environmental isolates, including four from Georgia agricultural sites. To verify that these mutations were associated with fungicide resistance, we tested growth on media amended with QoI or MBC fungicides for the four Georgia isolates carrying *cytB* and *benA* mutations (eAF222, eAF233, eAF234, eAF513) and four Georgia isolates not carrying these mutations (eAF77, eAF94, eAF128, eAF537). Only isolates with the *cytB* G143A mutation grew on medium with the QoI fungicide azoxystrobin and only isolates with the *benA* F219Y mutation grew on medium with the MBC fungicide benomyl (Supplementary Figure S2, Supplementary Table S4).

**TABLE 2.** Mutations associated with fungicide resistance in pan-azole-resistant *A. fumigatus*.

| Isolate | Source | *cyp51A* – azoles <sup>a</sup> | *cytB* – QoI <sup>b</sup> | *benA* – MBC <sup>c</sup> |
| --- | --- | --- | --- | --- |
| A1163 | Clinic | WT | WT | WT |
| eAF222 | Environment, 2018<br>USA | TR<sub>46</sub>/Y121F/T289A | G143A | F219Y |
| eAF233 | Environment, 2018<br>USA | TR<sub>46</sub>/Y121F/T289A | G143A | F219Y |
| eAF234 | Environment, 2018<br>USA | TR<sub>46</sub>/Y121F/T289A | G143A | F219Y |
| eAF513 | Environment, 2018<br>USA | TR<sub>46</sub>/Y121F/T289A | G143A | F219Y |
| 08-12-12-13 | Clinic, 2003<br>Netherlands | TR<sub>34</sub>/L98H/ | WT | F219Y |
| 08-31-08-91 | Clinic, 2004<br>Netherlands | TR<sub>34</sub>/L98H | WT | F219Y |
| 08-36-03-25 | Clinic, 2005<br>Netherlands | TR<sub>34</sub>/L98H | WT | F219Y |
| 10-01-02-27 | Clinic, 2010<br>Netherlands | TR<sub>34</sub>/L98H | G143A | F219Y |
| Afu 942/09 | Clinic, 2009<br>India | TR<sub>34</sub>/L98H | G143A | F219Y |
| Afu 1042/09 | Clinic, 2009<br>India | TR<sub>34</sub>/L98H | G143A | F219Y |
| Afu 124/E11 | Environment, 2011<br>India | TR<sub>34</sub>/L98H | G143A | F219Y |
| Afu 166/E11 | Environment, 2011<br>India | TR<sub>34</sub>/L98H | G143A | F219Y |
| Afu 218/E11 | Environment, 2011<br>India | TR<sub>34</sub>/L98H | G143A | F219Y |
| Afu 257/E11 | Environment, 2011<br>India | TR<sub>34</sub>/L98H | G143A | F219Y |
| Afu 343/P11 | Clinic, 2011<br>India | TR<sub>34</sub>/L98H | G143A | F219Y |
| Afu 591/12 | Clinic, 2012<br>India | TR<sub>34</sub>/L98H | G143A | F219Y |
| DI 15-96 | Clinic, 2008<br>USA | TR<sub>46</sub>/Y121F/T289A | G143A | F219Y |
| DI 15-102 | Clinic, 2010<br>USA | TR<sub>34</sub>/L98H | WT | F219Y |
| DI 15-106 | Clinic, 2012<br>USA | TR<sub>46</sub>/Y121F/T289A | G143A | F219Y |
| DI 15-116 | Clinic, 2014<br>USA | TR<sub>34</sub>/L98H | WT | F219Y |

<sup>a</sup> GenBank accession number EDP50065 from azole-sensitive isolate A1163 was used as wildtype for Cyp51A. Isolates 08-12-12-13 and 08-36-03-25 also carried S297T/F495I mutations for Cyp51A, but these have not been associated with azole resistance.
<sup>b</sup> GenBank accession number YP\_005353050 from azole-sensitive isolate A1163 was used as wildtype for CytB. All *cyp51A* TR mutants also carried V13I/I119V mutations for CytB, but these have not been associated with QoI resistance.
<sup>c</sup> GenBank accession number EDP56324 from azole-sensitive isolate A1163 was used as wildtype for BenA.

Fourteen of 19 pan-azole-resistant isolates included in our study also carried mutations for QoI resistance, MBC resistance, or both. All QoI-resistant and MBC-resistant mutants were also pan-azole resistant and clustered in the well-supported pan-azole clade (Figure 1, red branches). Eight of these multi-fungicide-resistant isolates were from agricultural sources in the United States and India and 6 were from clinical sources in the Netherlands and India. Four of these clinical isolates carried mutations conferring resistance to both MBC and QoI fungicides. That pan-azole-resistant *A. fumigatus* strains from patients carry the mutation for resistance to QoI fungicides used exclusively for plant protection shows definitively that these strains have an agricultural origin. However, not all pan-azole resistant isolates in our study showed this definitive signature of agricultural origin. Five of 19 pan-azole-resistant clinical isolates did not carry MBC or QoI fungicide-resistance mutations raising the possibility that they could have originated from nonagricultural sources. We found pan-azole-resistant *A. fumigatus* in a single clade on three continents. We also found resistance to MBC and QoI fungicides exclusively in this clade and always in combination with pan-azole resistance. Beyond showing the agricultural origin of clinical pan-azole resistance, our results suggest that the unique genetics of the pan-azole clade enable the evolution and/or persistence of antimicrobial resistance mutations.

## Methods

### Sampling

Between July 2017 and March 2018 we collected soil, plant debris, or compost from 56 agricultural sites in Georgia or Florida, USA, including 53 sites that had recently been treated with triazole fungicides, two sites that were in organic production with no triazole use in at least 10 years, and one compost pile with an unknown history of fungicide use on the plant material prior to composting (Supplementary Table S1). Each site was defined as a different field location, different crop at the same field location, or different triazole fungicide treatment. When soil was sampled, plant debris on the soil surface was included if available. If larger piles of debris were present on the soil surface they were collected separately. Soil was sampled by taking 5-10 soil cores to a depth of approximately 10 cm. Plant debris was sampled from the soil surface, cull piles at farms, and waste piles at pecan processing facilities. For each site we collected 4 samples at separate locations to minimize the collection of clones. Samples were stored in sealed plastic bags for transport and stored open to allow for gas exchange at 4°C for 2-20 d.

### Isolation and storage

The samples were processed as described previously by Snelder et al.^9^ and Hurst et al.^16^ with some modifications. Briefly, 2 g of soil was suspended in 15 ml of sterile 0.1 M sodium pyrophosphate. Samples were vortexed for 30 s and allowed to settle for 1 min. From the supernatant, 100 μl was pipetted onto Sabouraud dextrose agar (SDA) in 100-mm Petri dishes supplemented with 50 μg/ml chloramphenicol (Sigma Aldrich) and 5 μg/ml gentamicin (Research Products International). The dishes were incubated for 2 to 4 d at 45°C. Colonies of *A. fumigatus* were initially identified based on morphology and screened for azole resistance on SDA supplemented with 3 μg/ml tebuconazole (TEB; Bayer Corp). Many of the isolates that grew on 3 μg/ml TEB-amended solid medium were not able to grow at similar concentrations of TEB in liquid medium during the MIC testing described below; therefore, these isolates are designated as sensitive to TEB in Supplementary Table S1. Isolates were stored at −80°C in 15% glycerol.

### Antifungal susceptibility testing by Minimum Inhibitory Concentrations (MIC)

One hundred-seventy-two environmental *A. fumigatus* isolates (Supplementary Table S1), and 48 clinical isolates were tested for antifungal susceptibility under conditions described in the Clinical Laboratory Standard Institute broth microdilution method^29^. Antifungals tested included tebuconazole (TEB; TCI America, Oregon, USA), itraconazole (ITC; Thermo Sci Acros Organics, New Jersey, USA), voriconazole (VOR; Thermo Sci Acros Organics, New Jersey, USA), and posaconazole (POS; Apexbio Technology, Texas, USA) suspended in DMSO. Briefly, isolates were grown on complete media^30^ slants for 5 to 7 d at 37°C and harvested with 2 ml of 0.02% Tween-20 solution. Spore suspensions were adjusted to 0.09 – 0.13 OD at 530 nm using a spectrophotometer. One-hundred microliters of 2 ×10^4^ to 5 ×10^4^ conidia/ml were added to 100 ml of RPMI 1640 liquid medium (Thermo Sci Gibco, California, USA) in microtiter plates with final concentrations of antifungals ranging from 0.015 to 16 μg/ml. Plates were incubated at 37°C for 48 h and MIC break points were determined visually. MIC break point was defined as the lowest concentration at which there was 100% inhibition of growth. Assays were repeated for all resistant isolates and most sensitive isolates. For classification of sensitivity or resistance for ITC and VOR, we used the recommended clinical breakpoints of antifungal resistance for *A. fumigatus*^17^ which are > 2 μg/ml; however, EUCAST notes there is uncertainty regarding the cutoff values for POS and some data suggest > 1 μg/ml, which we use here, may be more relevant.

### DNA extraction

High molecular weight genomic DNA of *A. fumigatus* isolates was extracted using a modified CTAB protocol as described previously^31^. Briefly, approximately 100 mg of mycelium collected from cultures that had been incubated overnight in liquid complete medium^30^ were transferred to 2 ml tubes containing approximately 200 μl of 0.5-mm disruption glass beads (RPI, catalog #9831) and three 3-mm steel beads and lyophilized. Lyophilized cells were pulverized using Geno/Grinder at 1750 rpm for 30 s. Pulverized tissue was incubated in 1 ml of CTAB lysis buffer (100 mM Tris pH 8.0, 10 mM EDTA, 1% CTAB, 1% BME) for 30 min at 65°C. The samples were extracted with chloroform (500 μl) twice and DNA in the upper layer was precipitated in ice cold isopropanol. The precipitated DNA samples were washed with 70% ethanol twice, air dried, and dissolved in 100 μl sterile water. DNA was quantified using NanoDrop One (Thermo Sci, New Jersey, USA).

### cyp51A *sequencing*

For all environmental isolates in this study, *cyp51A* was PCR-amplified using previously designed primers^32^. PCR reactions were performed with the Q5 Hot Start High-Fidelity 2× Master Mix (New England Biolabs) with 100 ng of genomic DNA, 0.5 μM forward primer 5’-CGGGCTGGAGATACTATGGCT-3’ and 0.5 μM reverse primer 5’-GTATAATACACCTATTCCGATCACACC-3’ in 20 μl reactions. PCR reactions were performed at 98°C for 2 min followed by 30 cycles of 98°C for 15 s, 62°C for 15 s, and 72°C for 2:30 min, followed by a final extension at 72°C for 5 min. Amplicons were sequenced by the Sanger method (Eurofins genomics, USA) using primers 5’-GCATTCTGAAACACGTGCGTAG-3’, 5’-GTCTCCTCGAAATGGTGCCG-3’, and 5’-CGTTCCAAACTCACGGCTGA-3’. Promoter sequences were aligned to A1163 genomic sequence v43 from Ensembl. Coding sequences were translated to protein sequences and aligned to the Cyp51A protein of *A. fumigatus* A1163 (GenBank accession EDP50065). Sequence analysis was performed using Geneious v11.1.5 (Biomatters, Auckland, NZ).

### Microsatellite genotyping

Nine microsatellite markers (STR *Af* 2A, 2B, 2C, 3A, 3B, 3C, 4A, 4B, and 4C) previously developed for *A. fumigatus* (de Valk et al. 2005) were used to genotype 166 environmental isolates (Supplementary Table 1), the reference isolate Af293, and 48 clinical isolates from Georgia or Florida provided by the Mycotic Diseases Group at CDC^23^. Multiplex PCR was performed using the Type-it Microsatellite PCR kit (Qiagen) following the manufacturer’s protocol, but with the reaction volume modified to 10 μl. Multiplex reactions contained the following: 5 μl of 2× Type-it Master Mix, 1 μl of 10× primer mix (2 μM of each primer in the multiplex), 1 μl of DNA template, and RNAse-free water. Thermal cycling conditions had an initial denaturation at 95°C for 5 min followed by: 28 cycles of 95°C for 30 s, 57°C for 90 s, and 72°C for 30 s and a final elongation of 60°C for 30 min. Amplification of a product was confirmed by electrophoresis on a 1% (w/v) agarose gel with 1× TBE buffer. PCR products were diluted (1:15) then 1 μl of diluted PCR product was added with 0.1 μl of the internal size standard Genescan-500 Liz (Applied Biosystems) and 9.9 μl of Hi-Di formamide (Applied Biosystems). These were then incubated for 5 min at 95°C and placed immediately on ice. Fragment analysis was performed at the Georgia Genomics and Bioinformatics Core (GGBC) on an Applied Biosystems 3730×1 96-capillary DNA analyzer. Microsatellite Plugin in Geneious v.6 (Biomatters) was used to score alleles and loci were distinguished based on expected size range. To examine relationships among all isolates and isolates from different environments, a minimum spanning network was constructed using the Bruvo’s genetic distance model^33^ in the Poppr package executed in R^34^.

### Library preparation and whole genome sequencing

Genomic DNA was sheared to a mean size of 600 bp using a Covaris LE220 focused ultrasonicator (Covaris Inc., Woburn, MA). DNA fragments were Ampure (Beckman Coulter Inc., Indianapolis, IN) cleaned and used to prepare dual-indexed sequencing libraries using NEBNext Ultra library prep reagents (New England Biolabs Inc., Ipswich, MA) and barcoding indices synthesized in the CDC Biotechnology Core Facility. Libraries were analyzed for size and concentration, pooled and denatured for loading onto flowcell for cluster generation. Sequencing was performed on an Illumina Hiseq using 300×300 cycle paired-end sequencing kit. On completion, sequence reads were filtered for read quality, base called and demultiplexed using Casava v1.8.4. All raw reads and assemblies were deposited in GenBank under project #XXXXXX.

### SNP calling and neighbor-joining tree

Cleaned whole genome sequence reads for each isolate were *de novo* assembled using SPAdes v3.12.0^35^ with option --careful and trained to Af293 reference genome^36^ using option --trusted-contigs. Corrected fasta files generated from SPAdes were aligned to Af293 reference genome using Burrows-Wheeler Aligner (BWA) alignment tool^37^ and duplicate reads were marked using Picard v2.16.0. Single nucleotide polymorphisms (SNPs) were called with SAMtools mpileup v1.6^37^ with option -I to exclude insertions and deletions then with BCFtools v1.9 with option –c. Bases with phred quality score lower than 40 were filtered using SAMtools seqtk v1.2. SNP data were converted into interleaved phylip format and a neighbor-joining tree was constructed using Seaview v4.7^38^. Support for internal branches was determined by 100 bootstrap replicates. The tree was visualized and annotated using iTOL: International Tree of Life v5^39^.

### Genome mining for agricultural fungicide resistance

Whole genome sequences (Supplementary Table S3) were searched by tblastn^40^ for *A. fumigatus cyp51A* (GenBank accession number EDP50065), *benA* (GenBank accession number EDP56324), and *cytB* (GenBank accession number AFE02831). Blast hits were extracted from assemblies using BEDtools v2.26.0. Sequence analysis was performed using Geneious v11.1.5 (Biomatters, Auckland, NZ).

### Fungicide resistance phenotype assays

Sensitivity assays for QoI were conducted in 100-mm Petri dishes of Sabouraud dextrose agar (SDA) that contained 10 μg/ml of azoxystrobin (Sigma Aldrich analytical-grade, diluted from 10 mg/mL stock in acetone) and 100 μg/ml salicylhydroxamic acid (SHAM) (Sigma Aldrich analytical-grade, diluted from 100 mg/ml stock in methanol)^41^. Controls were identical except that they lacked azoxystrobin. Sensitivity assays for MBC fungicides were similar, except they contained only 10 μg/ml benomyl (Sigma Aldrich, diluted from10 mg/ml stock in DMSO) in SDA^42^. Controls lacked Benomyl. Three azoxystrobin-amended SDA and three control dishes, as well as 3 benomyl-amended and control dishes were inoculated with 100 μl of a 1 × 10^3^ conidia/ml *A. fumigatus* stock, spread using a sterile glass rod, and incubated at 37°C for 23 h at which point microcolonies were counted by eye. The experiment was performed twice.

## Acknowledgements

Research was funded by the Centers for Disease Control and Prevention (CDC; contract 200-2017-96199 to M.M. and M.T.B.) and the University of Georgia (President’s Interdisciplinary Seed Grant to M.T.B and M.M.). We thank the Mycotic Diseases Branch and Elizabeth Berkow at CDC for guidance and clinical isolates from Georgia and Florida. We thank Megan Dewdney, Gary Vallad, and Katia Xavier from the University of Florida, and Bhabesh Dutta, Robert Kemerait, Albert Culbreath, Tim Brenneman, Renee Allen, and Elizabeth Little from the University of Georgia for assistance collecting environmental samples. We thank Douda Bensasson from the University of Georgia for advice and insight on phylogenetic analysis.

## Author Contributions

S.E.K., M.T.B., and M.M. conceived and designed the research. S.E.K., T.M., L.G.S., and B.M. conducted the experiments. S.E.K., M.T.B., and L.G.S. collected environmental samples. S.E.K., T.M., L.G.S., B.M., M.M. and M.T.B analyzed the data. S.E.K., M.M. and M.T.B. wrote the manuscript with contributions from all authors.

## Materials requests & Correspondence

should be addressed to M.T.B. or M.M.

---

**SUPPLEMENTARY TABLE S1.**
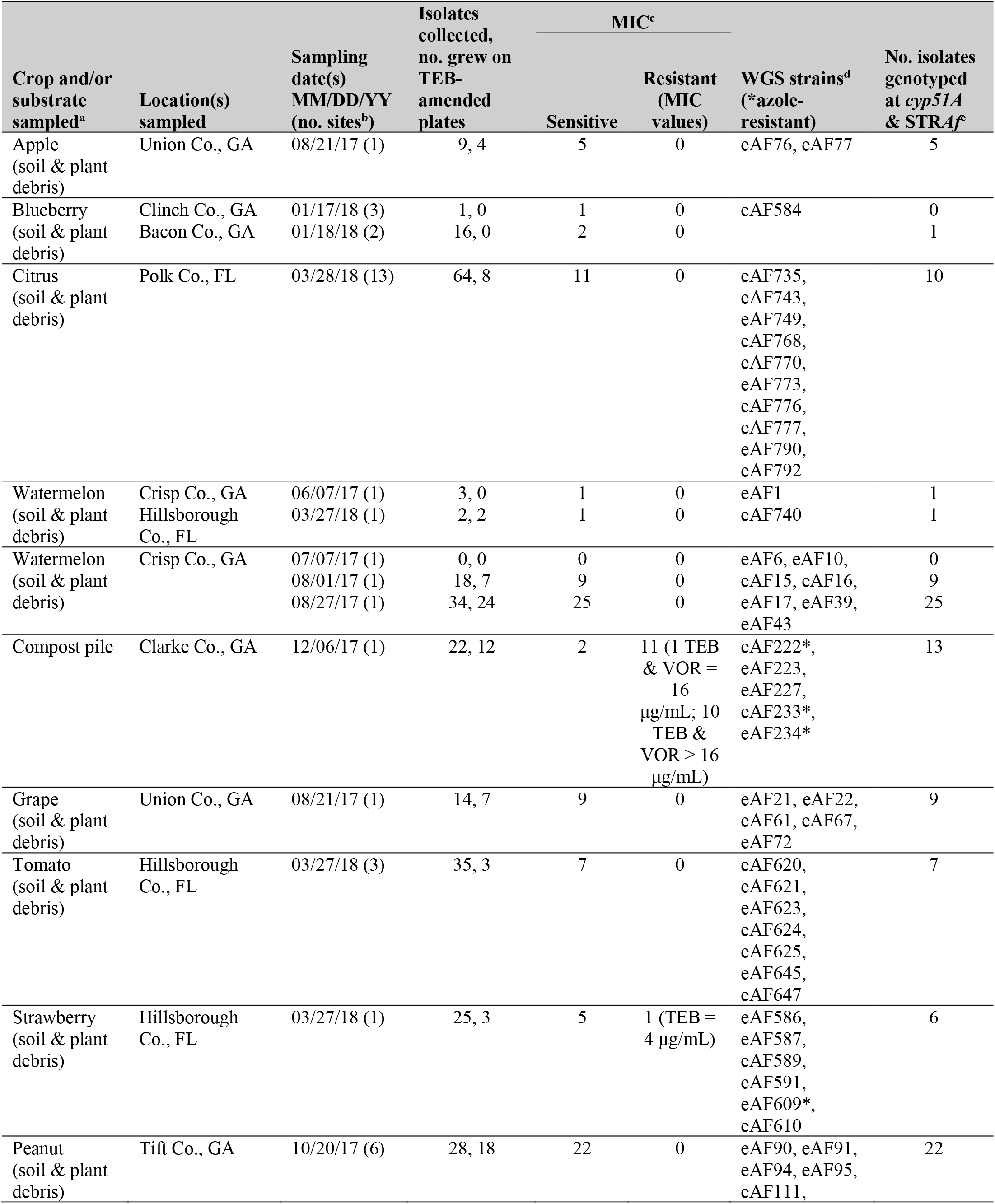
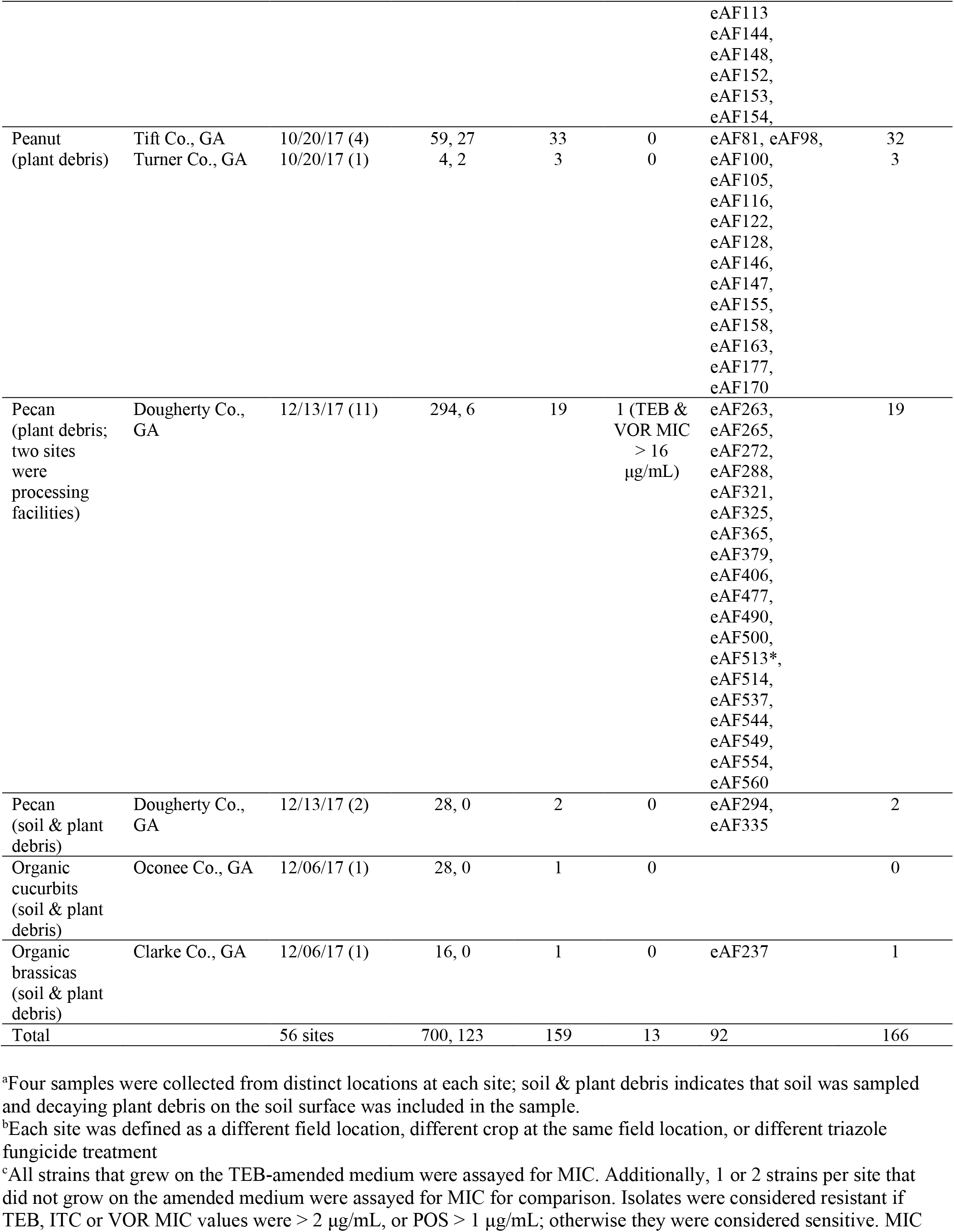
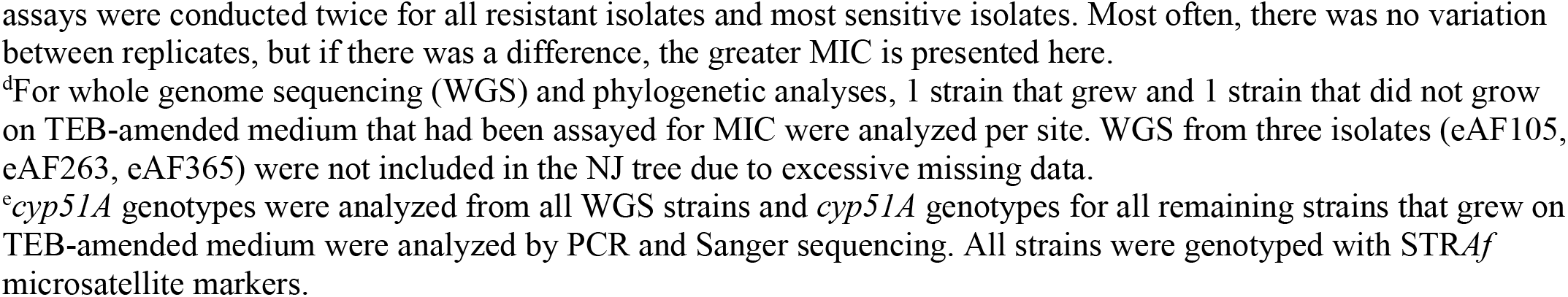
Sampling of *A. fumigatus* strains from agricultural sites.

**SUPPLEMENTARY TABLE S2.** Minimum inhibitory concentrations (MIC)^a^ for *A. fumigatus* (*n* = 172) isolated from agricultural environments in Georgia and Florida where azole fungicides were applied.

| Azole | Final Drug Concentration (µg/mL) |  |  |  |  |  |  |  |  |
| --- | --- | --- | --- | --- | --- | --- | --- | --- | --- |
|  | >16 | 16 | 8 | 4 | 2 | 1 | 0.5 | 0.25 | <0.25 |
| **Tebuconazole** | 11 | 1 |  | 1 | 85 | 68 | 6 |  |  |
| **Itraconazole** |  |  |  |  | 11 | 140 | 21 |  |  |
| **Voriconazole** | 12 |  |  |  |  | 1 | 81 | 72 | 6 |
| **Posaconazole** |  |  |  |  |  | 15 | 93 | 58 | 6 |

<sup>a</sup>Isolates were assayed for MIC once (some sensitive isolates) or twice (all resistant and some sensitive isolates). Most often, there was no variation between replicates, but if there was a difference, the greater MIC is presented here.

**SUPPLEMENTARY TABLE S3.** Publicly available genome sequence data used in this study for neighbor-joining tree and mining for antifungal-resistance genes and mutations (italics)

| NCBI_SRA_ID | Taxa_ID | Sampling Location | Environ (E) /Clinical (C) | Azole-Res (R) /Sens(S) | Reference |
| --- | --- | --- | --- | --- | --- |
| ERR769506 | 08-12-12-13 | Netherlands | C | R | doi: 10.1128/mBio.00536-15 |
| ERS663179 | 08-19-02-10 | Netherlands | E | R | doi: 10.1128/mBio.00536-15 |
| ERS663176 | 08-19-02-30 | Netherlands | E | S | doi: 10.1128/mBio.00536-15 |
| ERR769512 | 08-19-02-46 | Netherlands | E | R | doi: 10.1128/mBio.00536-15 |
| ERR769509 | 08-19-02-61 | Netherlands | E | R | doi: 10.1128/mBio.00536-15 |
| ERR769508 | 08-31-08-91 | Netherlands | C | R | doi: 10.1128/mBio.00536-15 |
| ERS663166 | 09-7500806 | UK | C | S | doi: 10.1128/mBio.00536-15 |
| ERR769511 | 10-01-02-27 | Netherlands | C | R | doi: 10.1128/mBio.00536-15 |
| ERS663168 | 12-7504462 | UK | C | S | doi: 10.1128/mBio.00536-15 |
| ERS663167 | 12-7504652 | UK | C | S | doi: 10.1128/mBio.00536-15 |
| ERS663169 | 12-7505054 | UK | C | S | doi: 10.1128/mBio.00536-15 |
| ERS663165 | 12-7505220 | UK | C | R | doi: 10.1128/mBio.00536-15 |
| ERS663164 | 12-7505446 | UK | C | R | doi: 10.1128/mBio.00536-15 |
| ERR232426 | A1163 | France | C | S | NCBI SRA |
| ERS663170 | Af293 | UK | C | S | doi: 10.1128/mBio.00536-15 |
| ERX207014 | AF300 | UK | C | S | NCBI SRA |
| SRS375791 | AF41 | UK | C | S | NCBI SRA |
| ERS663171 | AF65 | UK | C | S | doi: 10.1128/mBio.00536-15 |
| SRR617721 | AF72 | UK | C | R | doi: 10.1128/AAC.41.6.1364 |
| SRR617722 | AF90 | UK | E | R | doi: 10.1128/AAC.41.6.1364 |
| ERS663181 | Afu_1042/09 | India | C | R | doi: 10.1128/mBio.00536-15 |
| ERS663184 | Afu_124/E11 | India | E | R | doi: 10.1128/mBio.00536-15 |
| ERS663185 | Afu_166/E11 | India | E | R | doi: 10.1128/mBio.00536-15 |
| ERS663187 | Afu_218/E11 | India | E | R | doi: 10.1128/mBio.00536-15 |
| ERS663186 | Afu_257/E11 | India | E | R | doi: 10.1128/mBio.00536-15 |
| ERS663183 | Afu_591/12 | India | C | R | doi: 10.1128/mBio.00536-15 |
| ERS663180 | Afu_942/09 | India | C | R | doi: 10.1128/mBio.00536-15 |
| ERS216929 | AP65 | UK | E | S | NCBI SRA |
| SRR7418943 | ATCC_204305 | USA | C | S | doi: 10.3390/genes9070363 |
| SRR7418935 | ATCC_46645 | UK | C | S | doi: 10.3390/genes9070363 |
| ERS216946 | CF098 | UK | C | S | NCBI SRA |
| ERR232405 | CF337 | UK | C | R | NCBI SRA |
| SRR7418947 | CM2141 | Spain | C | S | doi: 10.3390/genes9070363 |
| SRR7418942 | CM237 | Spain | C | S | doi: 10.3390/genes9070363 |
| SRR7418945 | CM3248 | Spain | C | S | doi: 10.3390/genes9070363 |
| SRR7418944 | CM5419 | Spain | C | S | doi: 10.3390/genes9070363 |
| SRR7418949 | CM5757 | Spain | C | S | doi: 10.3390/genes9070363 |
| SRR7418936 | CM6458 | Spain | C | S | doi: 10.3390/genes9070363 |
| SRR7418938 | CM7555 | Spain | C | R | doi: 10.3390/genes9070363 |
| SRR7418941 | CM7632 | Spain | C | S | doi: 10.3390/genes9070363 |
| ERR232430 | D17 | UK | E | R | NCBI SRA |
| SRR617724 | F12219 | UK | C | R | doi: 10.3201/eid1507.090043 |
| SRR617725 | F12636 | UK | C | R | doi: 10.3201/eid1507.090043 |
| SRR617727 | F13619 | UK | C | R | doi: 10.3201/eid1507.090043 |
| SRR617729 | F14403 | UK | C | R | doi: 10.3201/eid1507.090043 |
| SRR617731 | F14532 | UK | C | R | doi:<br>10.3201/eid1507.090043 |
| SRS375803 | F14946G | UK | C | S | doi:<br>10.3201/eid1507.090043 |
| SRR617733 | F15390 | UK | C | R | doi:<br>10.3201/eid1507.090043 |
| SRR617735 | F16134 | UK | C | R | doi:<br>10.3201/eid1507.090043 |
| SRS375808 | F16311 | UK | C | S | doi:<br>10.3201/eid1507.090043 |
| ERR232417 | F17999 | UK | C | R | doi:<br>10.3201/eid1507.090043 |
| SRS375813 | F18085 | UK | C | S | doi:<br>10.3201/eid1507.090043 |
| ERR232439 | F18304 | UK | C | R | doi:<br>10.3201/eid1507.090043 |
| ERR232418 | F18329 | UK | C | R | doi:<br>10.3201/eid1507.090043 |
| ERR232413 | F20140 | UK | C | R | doi:<br>10.3201/eid1507.090043 |
| ERR232440 | F21572 | UK | C | R | doi:<br>10.3201/eid1507.090043 |
| ERR232414 | F21732 | UK | C | R | doi:<br>10.3201/eid1507.090043 |
| ERR232421 | F21857 | UK | C | R | doi:<br>10.3201/eid1507.090043 |
| ERS216942 | F4S1B | UK | E | S | NCBI SRA |
| SRS375812 | F5211G | UK | C | S | doi:<br>10.3201/eid1507.090043 |
| SRS375815 | F7763 | UK | C | S | doi:<br>10.3201/eid1507.090043 |
| SRR4002443 | ISSFT-021 | International<br>Space Station | E | S | doi:<br>10.1128/mSphere.00227-16 |
| ERR232441 | RSF2S8 | UK | E | R | NCBI SRA |
| ERS216922 | SF1S5 | UK | E | S | NCBI SRA |
| ERS216948 | SF1S6 | UK | E | S | NCBI SRA |
| ERS216938 | SF2S6 | UK | E | S | NCBI SRA |
| ERS216937 | SF3S1 | UK | E | S | NCBI SRA |
| ERS216930 | SF3S10 | UK | E | S | NCBI SRA |
| ERS216918 | SF4S10 | UK | E | S | NCBI SRA |
| *ERR769507* | *08-36-03-25* | *Netherlands* | *C* | *R* | *doi: 10.1128/mBio.00536-15* |
| *ERS663182* | *Afu_343/P/11* | *India* | *C* | *R* | *doi: 10.1128/mBio.00536-15* |
| *SRR7841978* | *DI_15-102* | *USA* | *C* | *R* | *doi: 10.1128/mBio.00437-20* |
| *SRR7841983* | *DI_15-106* | *USA* | *C* | *R* | *doi: 10.1128/mBio.00437-23* |
| *SRR7841993* | *DI_15-96* | *USA* | *C* | *R* | *doi: 10.1128/mBio.00437-36* |
| *SRR7842000* | *DI_15-116* | *USA* | *C* | *R* | *doi: 10.1128/mBio.00437-31* |

**SUPPLEMENTARY TABLE S4.** Fungicide resistance genotypes and growth phenotypes for agricultural isolates from Georgia and Florida.

| Isolate | *cyp51A*<br>genotype | Growth on<br>Azole<sup>a</sup> | *cytB*<br>genotype | Growth on<br>QoI<sup>b</sup> | *benA*<br>genotype | Growth on<br>MBC<sup>c</sup> |
| --- | --- | --- | --- | --- | --- | --- |
| eAF77 | WT | - | WT | - | WT | - |
| eAF94 | WT | - | WT | - | WT | - |
| eAF128 | WT | - | WT | - | WT | - |
| eAF537 | WT | - | WT | - | WT | - |
| eAF222 | TR<sub>46</sub>/Y121F/T289A | + | G143A | + | F219Y | + |
| eAF233 | TR<sub>46</sub>/Y121F/T289A | + | G143A | + | F219Y | + |
| eAF234 | TR<sub>46</sub>/Y121F/T289A | + | G143A | + | F219Y | + |
| eAF513 | TR<sub>46</sub>/Y121F/T289A | + | G143A | + | F219Y | + |

<sup>a</sup> One hundred conidia were inoculated to solid medium containing 3 µg/ml tebuconazole (TEB) and incubated at 37°C for 23-24 h. These isolates also had MICs ≥ 16 µg/ml in TEB liquid medium (Table 1).
<sup>b</sup> One hundred conidia were inoculated to solid medium containing 10 µg/ml azoxystobin and incubated at 37°C for 23-24 h.
<sup>c</sup> One hundred conidia were inoculated to solid medium containing 10 µg/ml benomyl and incubated at 37°C for 23-24 h.

**Supplementary Figure S1.**
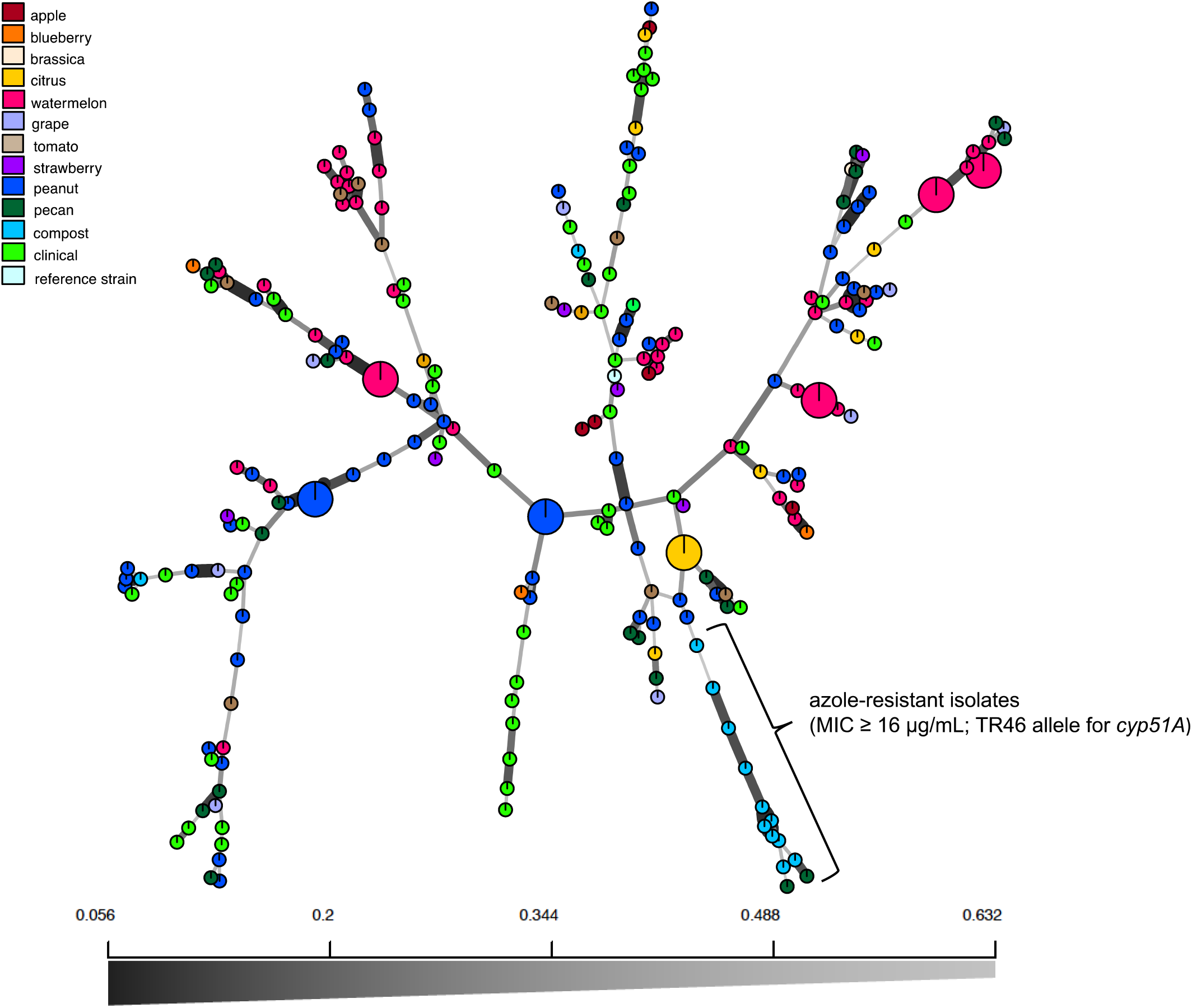
Minimum spanning network based on Bruvo’s genetic distance of agricultural and clinical isolates of *A. fumigatus* from Georgia and Florida. Isolates (168 agricultural and 48 clinical) were genotyped with 9 STR*Af* markers. Each circle represents a unique haplotype and the size of the circle represents the relative frequency of detection. The color of each circle represents the environment where the isolate was collected. Thicker lines represent shorter genetic distances. Individuals with the TR46/Y121F/T289A allele for *cyp51A* are shown in the lower right.

**Supplementary Figure S2.**
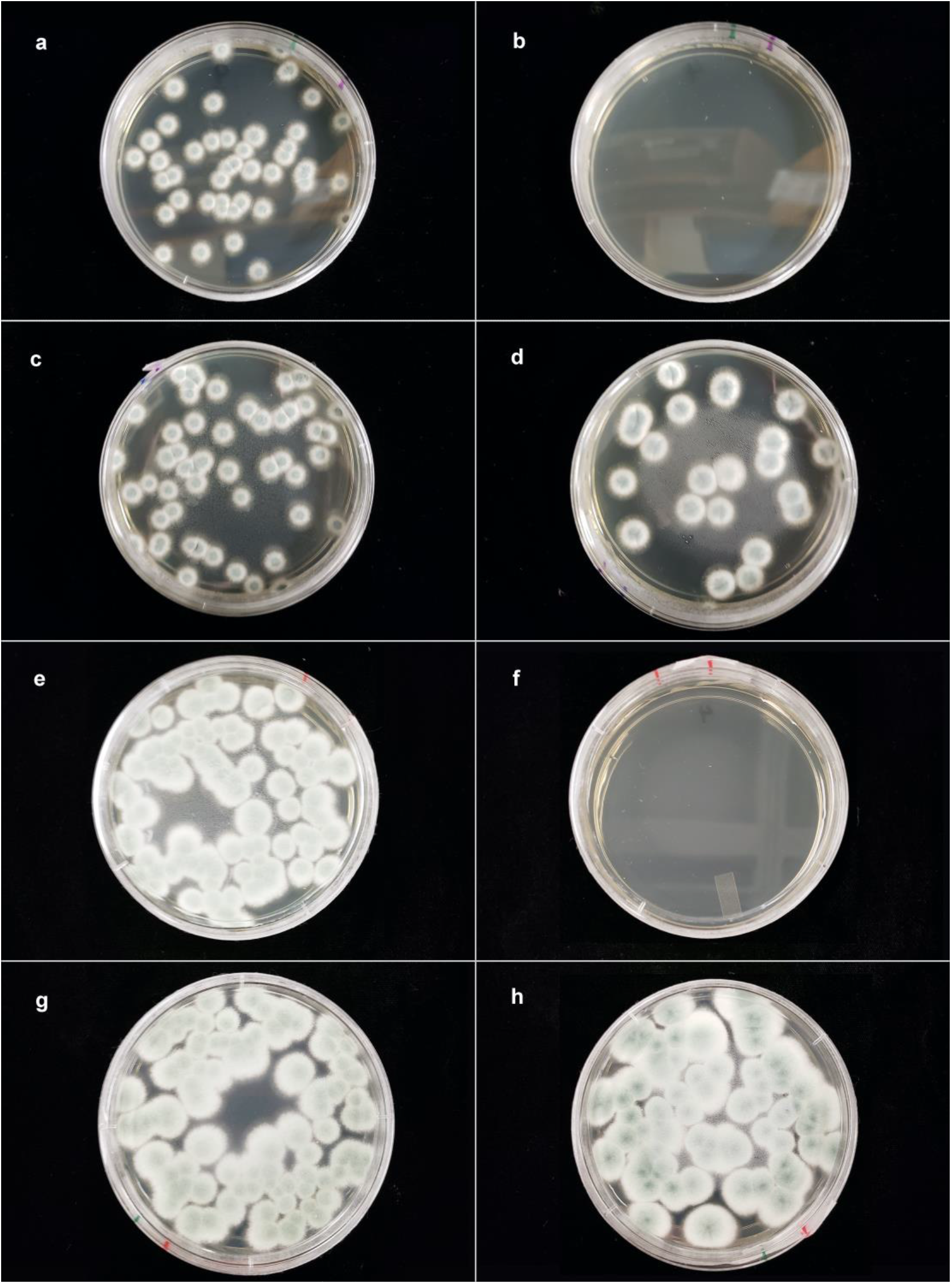
Pan-azole-resistant *A. fumigatus* (*cyp51A* TR_46_/Y121F/T289A) with *cytB* G143A and *benA* F219Y mutations are resistant to quinone outside inhibitor (QoI) and benzimidazole (MBC) fungicides. Left column (a, c, e, g) multi-fungicide-resistant isolate eAF222. Right column (b, d, f, h) sensitive isolate eAF94. (a, b) SDA (Sabouraud dextrose agar) + the QoI fungicide azoxystrobin + salicylhydroxamic acid (SHAM). (c, d) SDA medium + SHAM. (e,f) SDA medium + the MBC fungicide benomyl. (g, h) SDA medium.

